## Supplementary Information for "Structural basis for kinase inhibition in the tripartite *E. coli* HipBST toxin-antitoxin system"

SUPPLEMENTARY DATA

#### TABLE S1. Crystallographic data statistics

| | **HipBST$^{D233Q}$** | **HipBST$^{S57A}$** | **HipBST$^{S59A}$** |
|---|---|---|---|
| ***Data Collection*** | | | |
| Wavelength (Å) | 1.54180 | 0.97625 | 1.54180 |
| Resolution range (Å) | 49.9–2.9 (3.0–2.9)* | 46.9–2.4 (2.5–2.4)* | 50.4–3.34 (3.5 – 3.4)* |
| Space group | C121 | C121 | C121 |
| Unit cell dimensions | | | |
| a, b, c, (Å) | 281.66,106.47, 57.75 | 281.68, 106.07, 57.56 | 285.26, 107.15, 58.45 |
| *α, β, γ* (º) | 90, 90.75, 90 | 90, 90.65, 90 | 90, 90.71, 90 |
| Total reflections | 74,587 (7,335) | 625,868 (25,597) | 41,777 (4,068) |
| Unique reflections | 37,716 (3,719) | 66,117 (6,503) | 22,580 (2,193) |
| Multiplicity | 2.0 (2.0) | 9.5 (3.9) | 1.9 (1.9) |
| Completeness (%) | 99.8 (99.3) | 99.9 (99.4) | 87.71 (84.84) |
| $R_{meas}$ (%) | 0.13 (1.105) | 0.27 (0.08) | 0.17 (1.01) |
| I/σ(I) | 6.62 (0.86) | 10.3 (0.72) | 7.10 (0.93) |
| $CC_{1/2}$ | 0.99 (0.37) | 0.99 (0.37) | 0.97 (0.43) |
| ***Refinement*** | | | |
| Average B-factor (Å$^2$) | 98.5 | 86.1 | 119.4 |
| protein | 98.9 | 86.5 | 119.9 |
| ligands | - | 166.7 | 118.6 |
| solvent | 80.0 | 76.7 | 78.3 |
| No. of reflections | 37,706 (3,719) | 66,112 (6,504) | 22,570 (2,193) |
| No. of reflections (free) | 1,914 (175) | 3,364 (326) | 1086 (86) |
| R-work (%) | 19.5 (31.8) | 21.0 (42.8) | 20.2 (29.2) |
| R-free (%) | 22.8 (34.5) | 23.8 (46.2) | 24.3 (35.4) |
| Number of | | | |
| protein (residues) | 1,018 | 1,017 | 1,023 |
| solvent (atoms) | 203 | 402 | 110 |
| ligand (atoms) | - | 20 | 10 |
| rmsd (bonds, Å) | 0.011 | 0.013 | 0.013 |
| rmsd (angles, degrees) | 1.49 | 1.75 | 1.66 |
| Rotamer outliers (%) | 9.8 | 6.0 | 10.07 |
| Clashscore | 4.4 | 2.7 | 5.8 |
| Ramachandran statistics | | | |
| favoured (%) | 95.1 | 96.0 | 95.6 |
| allowed (%) | 4.7 | 3.6 | 3.9 |
| outliers (%) | 0.2 | 0.4 | 0.5 |
| Rama-Z score | | | |
| whole | -1.63 | -0.44 | -0.90 |
| helix | -1.27 | -0.40 | -0.48 |
| sheet | -1.36 | 0.21 | 0.07 |
| loop | -0.75 | -0.16 | -0.73 |

*Numbers in parentheses refer to the outermost resolution shell.

### TABLE S2. Bacterial strains and plasmids.

| *E. coli* strains | Description | Reference or source |
|---|---|---|
| **MG1655** | Wild-type K12 | (Blattner et al., 1997) |
| **TB28** | MG1655 Δ*lacIZYA* | Laboratory collection |

| Plasmid | Description[a] | Reference or source |
|---|---|---|
| **pBAD33** | p15 *araC* $P_{BAD}$, $Cm^r$ | (Guzman et al., 1995) |
| **pGH254** | Mini-R1, *lacZYA* transcriptional fusion vector, $Kan^r$ | Laboratory collection |
| **pNDM220** | Mini-R1 *lacI*$^q$ $P_{A1/04/03}$, $Amp^r$ | (Gotfredsen and Gerdes, 1998) |
| **pET-15b** | pBR322 *lacI* $P_{T7}$, $Amp^r$ | Novagen |
| **pKG127** | pUC57::*hipBST*$_{O127}$ | (Vang Nielsen et al., 2019) |
| **pSVN1** | pBAD33::*hipT*, start codon GTG | (Vang Nielsen et al., 2019) |
| **pSVN68** | pUC57::*hipB-S-T*$^{S57A}_{His6}$, optimized SDs for all genes | This work |
| **pSVN78** | pET-15b::*hipB-S-T*$^{S57A}_{His6}$, optimized SDs for all genes | This work |
| **pSVN88** | pUC57::*hipB-S-T*$^{D233Q}_{His6}$, optimized SDs for all genes | This work |
| **pSVN96** | pET-15b::*hipB-S-T*$^{D233Q}_{His6}$, optimized SDs for all genes | This work |
| **pSVN109** | pNDM220::*hipS*, optimized SD | (Vang Nielsen *et al.*, 2019) |
| **PSVN141** | pGH254::$P_{hipBST}$-*hipB*'-*laZ*, transcriptional $P_{hipBST}$-*hipB*' *lacZ* fusion | This work |
| **pSVN178** | pNDM220::*hipS*$^{W65A}$, optimized SD | This work |
| **pSVN181** | pBAD33::*hipB-S-T*$^{D233Q}$, optimized SDs for all genes | This work |
| **pSVN182** | pBAD33::*hipB-S*, optimized SDs for both genes | This work |
| **pSVN185** | pBAD33::*hipB-T*$^{D233Q}$, optimized SDs for both genes | This work |
| **pSVN188** | pBAD33::*hipS-T*$^{D233Q}$, optimized SDs for both genes | This work |
| **pSVN189** | pBAD33::*hipB*, optimized SD | This work |
| **pSVN190** | pBAD33::*hipS*, optimized SD | This work |
| **pSVN193** | pBAD33::*hipT*$^{D233Q}$, optimized SD | This work |
| **pSVN194** | pBAD33::*hipT*$^{S57D}$, start codon GTG | This work |
| **pSVN195** | pBAD33::*hipT*$^{S59D}$, start codon GTG | This work |
| **pSVN199** | pBAD33::*hipT*$^{S57A}$, start codon GTG | This work |
| **pSVN201** | pBAD33::*hipT*$^{S59A}$, start codon GTG | This work |
| **pSNN1** | pET-15b::*hipT*$^{S57A}_{His6}$, optimized SD | This work |
| **pSNN2** | pET-15b::*hipT*$^{S57A+D210A}_{His6}$, optimized SD | This work |
| **pMME3** | pET-15b::*hipB-S-T*$^{S59A}_{His6}$, optimized SDs for all genes | This work |
| **pRBS1** | pET-15b::*hipB-S-T*$^{D210A}_{His6}$, optimized SDs for all genes | This work |
| **pRBS2** | pET-15b::*hipB-S-T*$^{S57D,D210A}_{His6}$, optimized SDs for all genes | This work |
| **pRBS3** | pET-15b::*hipB-S-T*$^{S59D,D210A}_{His6}$, optimized SDs for all genes | This work |

[a]SD, Shine-Dalgarno sequence.

#### TABLE S3. Oligonucleotides and primers.

| Oligonucleotide | Sequence |
|---|---|
| **FP1(GTG)** | CCCCGTCGACGGATCCAAGGAGTTTTATAAGTGGCGAATTGTCGTATTCTG |
| **FP21** | GGGGGGTACCGGATCCAAAATAAGGAGGAAAAAAAAATGATCTGCTCAGGACCAC |
| **FP22** | CCCCCTCGAGGGATCCAAAATAAGGAGGAAAAAAAAATGCATCGGCGAGTGAAAG |
| **FP43** | CCCCGAATTCCTCTCCCGATGAGATCAGC |
| **FP46** | GGGGGTCGACCTGCAGAAAATAAGGAGGAAAAAAAAATGGCGAATTGTCGTATTCTG |
| **FP47** | GGGGGGTACCGGATCCAAAATAAGGAGGAAAAAAAAATGGCGAATTGTCGTATTCTG |
| **FP48** | GGGGGTCGACCTGCAGAAAATAAGGAGGAAAAAAAAATGGCGAATTGTCGTATTCTG |
| **RP1** | CCCCCGCATGCGAATTCGCTCACAGCAGCCCCAGACG |
| **RP11** | CCCCCTCGAGAAGCTTTCACAGCAGCCCCAGACG |
| **RP14** | GGGGGAATTCAAGCTTTTATTCCTCCCAAGGTAAAATC |
| **RP15** | GGGGGAATTCAAGCTTTCACTCGCCGATGCATAG |
| **RP32** | CCCCGGATCCTCTGCAACTCCTGGAGTTG |
| **RP42** | GGGGGTCGACCTGCAGTCACTCGCCGATGCATAG |
| **HipT S57D Fw** | GCGTCAACAAAAAGGGATGGATATTTCCGGTT |
| **HipT S57D Rv** | GGGCTGGTAACCGGAAATATCCATCCCTTTTT |
| **HipT S59D Fw** | GTCAACAAAAAGGGATGAGTATTGACGGTTAC |
| **HipT S59D Rv** | TTGGGCTGGTAACCGTCAATACTCATCCCTTT |
| **HipT S59A Fw** | GTCAACAAAAAGGGATGAGTATTGCCGGTTAC |
| **HipT S59A Rv** | TTGGGCTGGTAACCGGCAATACTCATCCCTTT |
| **hipT D210A Fw** | TAAATGCATCGCGTTATTACCCAGCAACAA |
| **hipT D210A Rv** | CTGGGTAATAACGCGATGCATTTACGAAACTTT |
| **hipT S57S59A Fw** | GGGATGAGTATTGCCGGTTACCAGCCCAAATTGCAA |
| **hipT S57S59A Rv** | GTAACCGGCAATACTCATCCCTTTTTGTTGACGCGG |
| **hipS W65A Fw** | CAGAAGGAGCTCTGCGTCAACGCTA |
| **hipS W65A Rv** | TGACGCAGAGCTCCTTCTGGCGC |
| **hipX S57A Fw** | AAGGGATGGCTATTTCCGGTTACCAGCC |
| **hipX S57A Rv** | CGGAAATAGCCATCCTTTTTGTTGACG |
| **hipX D233Q Fw** | CGGTGTATCAGTTTGTTTCTGTCGCTCCC |
| **hipX D233Q Rv** | GAAACAAACTGATACACCGGCGCTAACG |
| **hipBS del Fw** | ACGACAATTCGCCATTTTTTTTTCCTCCTTATTTTTCTAGAGGG |
| **hipBS del Rv** | TTCCCCTCTAGAAAAATAAGGAGGAAAAAAAAATGGCGAAT |
| **Q5 HipT D210A Fw** | GGTAATAACGctATGCATTTACGAAACTTTG |
| **Q5 HipT D210A Rv** | CAGCAACCAGGCGTAAAC |
| **Q5 HipT S57D Fw** | AAAGGGATGGaTATTTCCGGT |
| **Q5 HipT S57D Rv** | TTGTTGACGCGGAAGTTC |
| **Q5 HipT S59D Fw** | TAGACATCCCTTTTTGTTGACG |
| **Q5 HipT S59D Rv** | TTGATGGTTACCAGCCCAAATTG |

#### SUPPLEMENTARY METHODS

##### Construction of plasmids.

**pSVN68.** The S57A mutation in *hipB-S-T*$_{\text{His6}}$ with optimized Shine Dalgarno (SD) sequences for all three genes was created using pSVN61 (Vang Nielsen *et al.*, 2019) and primers hipX S57A Fw and hipX S57A Rv in a site-directed plasmid mutagenesis PCR. The samples were pooled and digested with DpnI to remove the template plasmid before being transformed into *E. coli* DH5α.

**pSVN78.** *hipB-S-T*$^{\text{S57A}}_{\text{His6}}$ with optimized SDs for all three genes was sub-cloned from pSVN68 by digesting with XbaI and XhoI, purifying the DNA fragment and ligating into pET-15b.

**pSVN88.** The D233Q mutation in *hipB-S-T*$_{\text{His6}}$ with optimized SD sequences for all three genes was created using pSVN61 (Vang Nielsen *et al.*, 2019) and primers hipX D233Q Fw and hipX D233Q Rv in a site-directed plasmid mutagenesis PCR. The samples were pooled and digested with DpnI to remove the template plasmid before being transformed into *E. coli* DH5α.

**pSVN96.** *hipB-S-T*$^{\text{D233Q}}_{\text{His6}}$ with optimized SDs for all three genes was sub-cloned from pSVN88 by digesting with XbaI and XhoI, purifying the DNA fragment and ligating into pET-15b.

**pSVN141.** pGH254::P$_{hipBST}$-*hipB*'-*laZ*, P$_{hipBST}$-*hipB*', a fragment containing 224 bp upstream of the *hipB* gene plus the first 73 bp of the *hipB* gene was amplified from pKG127 using primers FP43 and RP32. The resulting PCR product was digested with EcoRI and BamHI and ligated into pGH254.

**pSVN178.** *hipS*$^{\text{W65A}}$ was created by a two-step PCR reaction. Two fragments were amplified from pSVN109 using primers FP22 and *hipS* W65A Rv in one reaction and *hipS* W65A Fw and RP14 in the other. The resulting two PCR products were joined by a second round of PCR using both fragments as template DNA and primers FP22 and RP14. The final PCR product was digested with XhoI and EcoRI and ligated into pNDM220.

**pSVN181.** *hipB-S-T*$^{D233Q}$ with optimized SDs for all three genes was amplified from pSVN88 using primers FP21 and RP11. The resulting PCR product was digested with KpnI and HindIII and ligated into pBAD33.

**pSVN182.** *hipB-S* with optimized SDs for both genes was amplified from pSVN87 using primers FP21 and RP14. The resulting PCR product was digested with KpnI and HindIII and ligated into pBAD33.

**pSVN185.** *hipB-T*$^{D233Q}$ with optimized SDs for both genes was created by amplifying two PCR products from pSVN88: *hipB* using primers FP21 and RP42 and *hipT*$^{D233Q}$ using primers FP48 and RP11. The *hipB* fragment was digested with KpnI and PstI, while the *hipT*$^{D233Q}$ fragment was digested with PstI and HindIII. The two digested fragments were then ligated into pBAD33 using the KpnI and HindIII restriction sites.

**pSVN188.** *hipS-T*$^{D233Q}$ with optimized SDs for both genes was amplified from pSVN87 using primers FP46 and RP11. The resulting PCR product was digested with KpnI and HindIII and ligated into pBAD33.

**pSVN189.** *hipB* with optimized SD was amplified from pSVN87 using primers FP21 and RP15. The resulting PCR product was digested with KpnI and HindIII and ligated into pBAD33.

**pSVN190.** *hipS* with optimized SD was amplified from pSVN87 using primers FP46 and RP14. The resulting PCR product was digested with KpnI and HindIII and ligated into pBAD33.

**pSVN193.** *hipT*$^{D233Q}$ with optimized SD was amplified from pSVN88 using primers FP47 and RP11. The resulting PCR product was digested with KpnI and HindIII and ligated into pBAD33.

**pSVN194.** *hipT*$^{S57D}$ with start codon GTG was created by a two-step PCR reaction. Two fragments were amplified from pKG127 using primers FP1(GTG) and HipT S57D Rv in one reaction and HipT S57D Fw and RP1 in the other. The resulting two PCR products were joined by a second round of PCR using both fragments as template DNA and primers FP1(GTG) and RP1. The final PCR product was digested with SalI and SphI and ligated into pBAD33.

**pSVN195.** *hipT*$^{S59D}$ with start codon GTG was created by a two-step PCR reaction. Two fragments were amplified from pKG127 using primers FP1(GTG) and HipT S59D Rv in one reaction and HipT S59D Fw and RP1 in the other. The resulting two PCR products were joined by a second round of PCR using both fragments as template DNA and primers FP1(GTG) and RP1. The final PCR product was digested with SalI and SphI and ligated into pBAD33.

**pSVN199.** *hipT*$^{S57A}$ with start codon GTG was amplified from pSVN78 using primers FP1(GTG) and RP1. The resulting PCR product was digested with SalI and SphI and ligated into pBAD33.

**pSVN201.** *hipT*$^{S59A}$ with start codon GTG was created by a two-step PCR reaction. Two fragments were amplified from pKG127 using primers FP1(GTG) and HipT S59A Rv in one reaction and HipT S59A Fw and RP1 in the other. The resulting two PCR products were joined by a second round of PCR using both fragments as template DNA and primers FP1(GTG) and RP1. The final PCR product was digested with SalI and SphI and ligated into pBAD33.

**pSNN1**. *hipT*$^{S57A}$ with optimized SD sequence was amplified from pSVN78 using primers hipBS del Fw and hipBS del Rv. This resulted in the deletion of *hipB* and *hipS*.

**pSNN2.** *hipT*$^{S57A+D210A}$ with optimized SD sequence was amplified from pSNN1 using primers hipT D210A Fw and hipT D210A Rv.

**pMME3**. *hipBST*$^{S59A}$ with optimized SD sequence was amplified from pSVN78 using primers hipT S57S59A Fw and hipT S57S59A Rv to introduce the mutations A57S and S59A.

**pRBS1.** *hipB-S-T*$^{D210A}{}_{His6}$ in pET-15b was created using the Q5 Site-Directed Mutagenesis kit (NEB) according to the procedures from the manufacturer, using pSVN78 as template and primer Q5 HipT D210A Fw and Q5 HipT D210A Rv.

**pRBS2.** *hipB-S-T*$^{S57D,D210A}{}_{His6}$ in pET-15b was created using Q5 Site-Directed Mutagenesis kit (NEB) according to the procedures from the manufacturer, using pRBS1 as template and primer Q5 HipT S57D Fw and Q5 HipT S57D Rv.

**pRBS3.** *hipB-S-T*$^{S59D,D210A}{}_{His6}$ in pET-15b was created using Q5 Site-Directed Mutagenesis kit (NEB) according to the procedures from the manufacturer, using pRBS1 as template and primer Q5 HipT S59D Fw and Q5 HipT S59D Rv.

#### SUPPLEMENTARY FIGURES

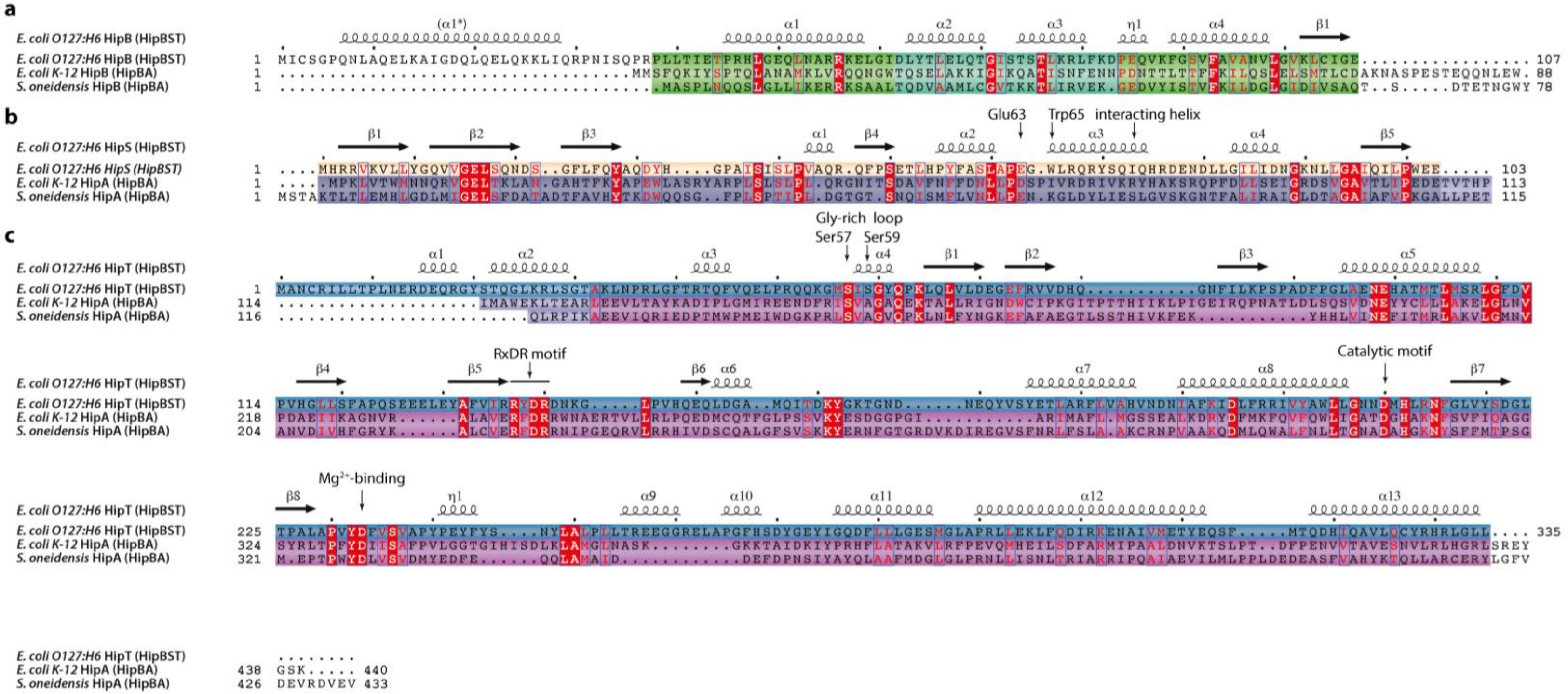


**Figure 1 supplement 1. Sequence alignment with consensus elements of HipBST, HipBA$_{Ec}$ and HipBA$_{So}$. a.** Alignment of *E. coli* O127:H6 HipB with HipB from *E. coli* K-12 HipBA and HipB from *S. oneidensis* HipBA. The secondary structure motifs observed in *E. coli* O127:H6 HipB is shown above the sequences, except for α1*, which is missing from the crystal structures but predicted by both JPred4 (Drozdetskiy *et al.*, 2015) and AlphaFold2 (Jumper *et al.*, 2021). **b.** Alignment of *E. coli* O127:H6 HipS with the N-subdomain 1 from both *E. coli* K-12 HipA and *S. oneidensis* HipA. The position of the Gly-rich loop interacting helix (α3, "interacting helix") and Glu63 and Trp65 of HipS are indicated. The secondary structure observed in *E. coli* O127:H6 HipS is shown above the sequences **c.** Alignment of *E. coli* O127:H6 HipT with the main kinase domains of *E. coli* K-12 HipA and *S. oneidensis* HipA. The position of the Gly-rich loop (residue 58-63), phosphorylation sites (Ser57 and Ser59), RxDR motif, catalytic motif (Asp210), and $Mg^{2+}$-binding motif (Asp233), are indicated. The secondary structure observed in *E. coli* O127:H6 HipT is shown above the sequences.

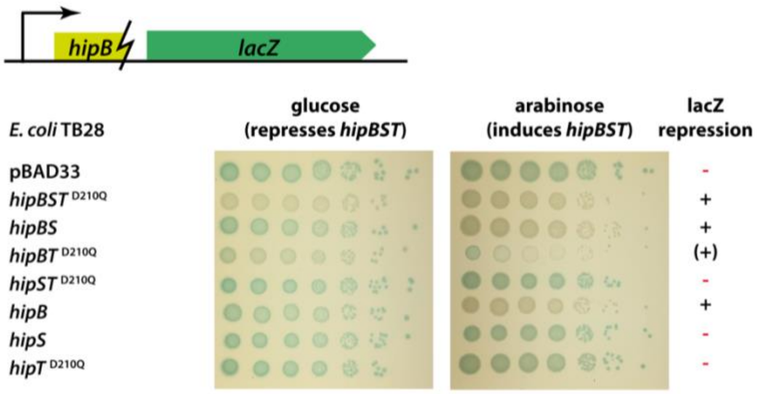


**Figure 1 supplement 2. Transcriptional regulation of the HipBST promoter of *E. coli* O127:H6.** All strains shown in this assay carried the reporter plasmid (pSVN141, pGH254::$P_{hipBST}$-*hipB'*-*lacZ*, top drawing) with the *hipBST* promoter region and a 5' fragment of the *hipB* gene (including 224 bp upstream of *hipB* plus the first 73 bp of the *hipB* orf) transcriptionally fused (indicated with a lightening symbol) to *lacZ*. The assay was made in strain *E. coli* TB28 harbouring the reporter plasmid and either empty pBAD33 vector or arabinose inducible combinations of *hipB*, *hipS*, and *hipT* in the context of the HipTD233Q inactive mutant. The strains were grown in liquid YT, diluted, and spotted onto YT agar plates containing 40 µg/ml X-gal (indicator turning blue upon *lacZ* expression) and 0.2% glucose (to repress *hipB/S/T*D233Q) or 0.2% arabinose (to induce *hipB/S/T*D233Q). Results are representative of two independent experiment and plasmid used were pSVN181 (pBAD33::*hipB*-*S*-*T*D233Q), pSVN182 (pBAD33::*hipB*-*S*), pSVN185 (pBAD33::*hipB*-*T*D233Q), pSVN188 (pBAD33::*hipS*-*T*D233Q), pSVN189 (pBAD33::*hipB*), pSVN190 (pBAD33::*hipS*), or pSVN193 (pBAD33::*hipT*D233Q).

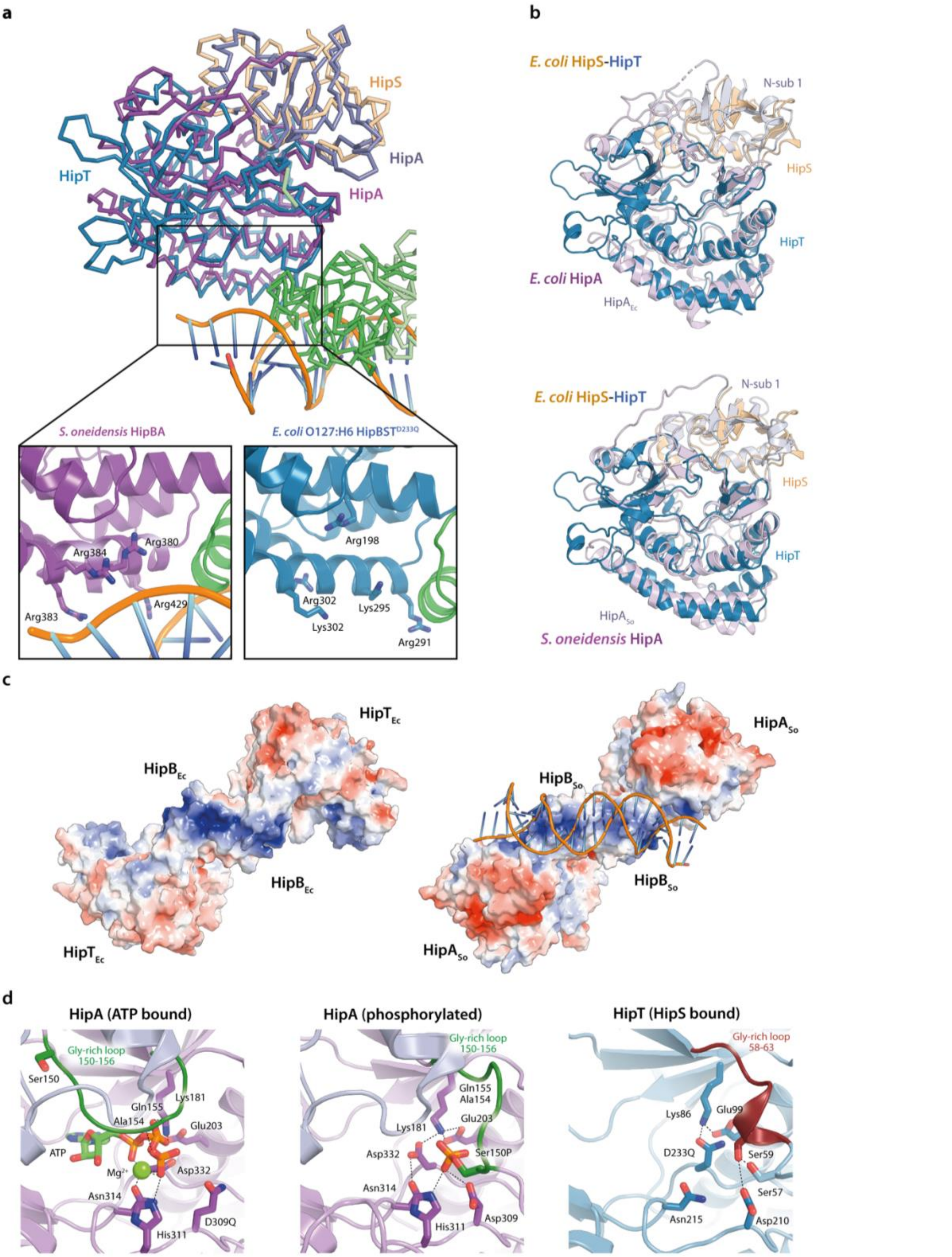


**Figure 1 supplement 3. Structural homolog analysis of HipBST to HipBA a.** Structural alignment of *S. oneidensis* HipBA in complex with DNA (PDB: 4PU3) with *E. coli* O127:H6 HipBST (this study) (Wen *et al.*, 2014). The close-up views highlight several positively charged residues in the area responsible for phosphate backbone interactions in *S. oneidensis* HipBA (left) and their counterparts in *E. coli* HipBST (right). **b.** Structural alignment of the *E. coli* O127:H6 HipST heterodimer with *E. coli* K-12 HipA (top) and *S. oneidensis* HipA (bottom). Protein domains and chains are indicated with labels in corresponding colours. **c.** APBS electrostatics of HipBST (left) and $HipBA_{So}$ in complex with DNA. The positive groove on the HipB-dimer is consistent between the two homologs. **d.** Close-up of the active site of *E. coli* K-12 HipA in the ATP-bound state (left), the same site in the Ser150 phosphorylated state (middle) and the corresponding region in HipT D233Q (right).

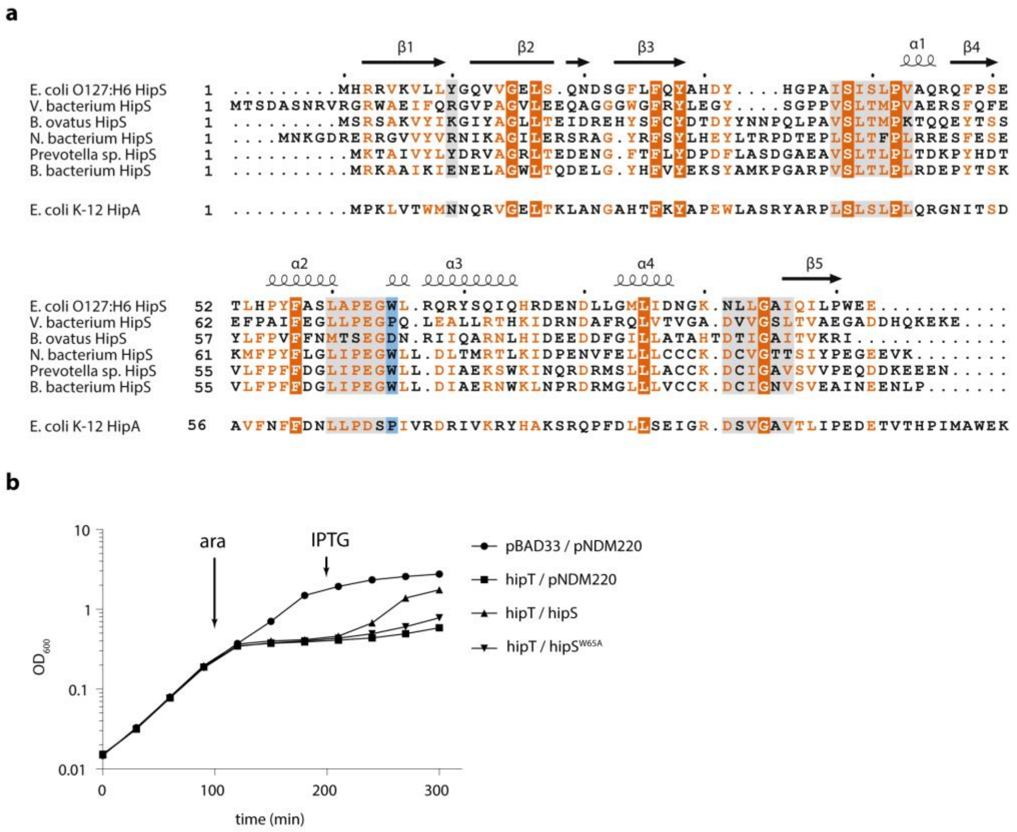


**Figure 2 supplement 1. Sequence and function of *E. coli* HipS. a.** Sequence alignment of *E. coli* O127:H6 HipS and selected orthologues with the N-subdomain 1 of *E. coli* K-12 HipA (bottom). The secondary structure observed for *E. coli* O127:H6 HipS is shown above the alignment. Fully conserved residues are in white text on a dark orange background, partially conserved residues in orange text, regions that interact with HipT are shown on a light grey background, and Trp65 that intercalates in HipT is shown on a blue background. **b.** Growth curves ($OD_{600}$) of *E. coli* MG1655 grown in YT medium and harbouring empty pBAD33 vector (pBAD33) or pSVN1 (pBAD33::*hipT*, "*hipT*"), in combination with empty pNDM220 vector (pNDM220), pSVN109 (pNDM220::*hipS*, "*hipS*"), or pSVN178 (pNDM220::*hipS*$^{W65A}$, *hipS*$^{W65A}$) as indicated. At the indicated times, 0.2% arabinose was added to induce *hipT* (ara) and 200 µM IPTG to induce *hipS* or *hipS*$^{W65A}$ (IPTG). The data points represent mean values of results from at least three independent experiments, and error bars show standard deviations (hidden when small).

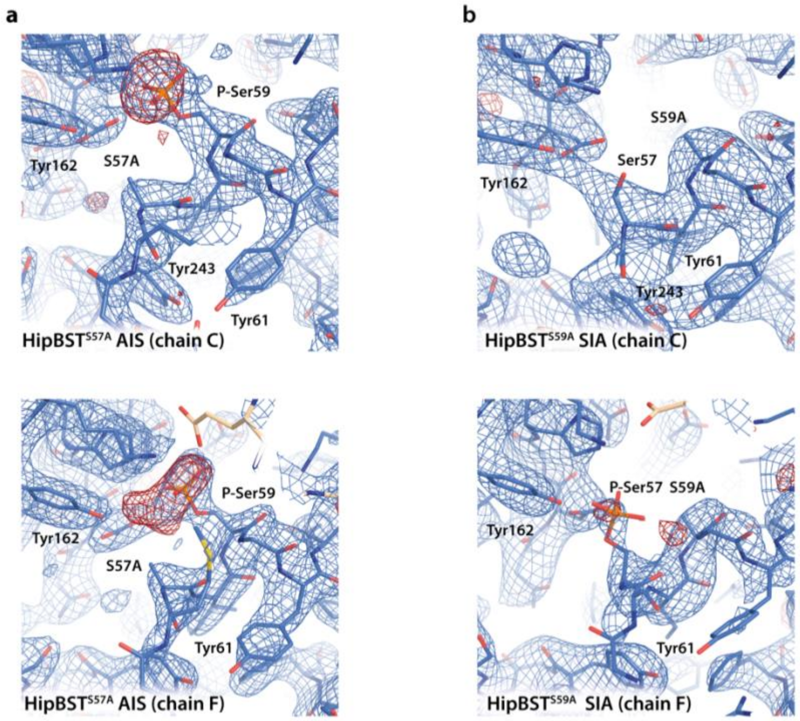


**Figure 3 supplement 1. Electron density surrounding the phosphoserine sites of HipT before and after modelling. a.** Electron density maps for the HipBST$^{S57A}$ structure showing the active sites of HipT in both copies in the ASU (chain C, top and chain F, bottom). In both cases, the $2F_o$-$F_c$ map (blue) contoured at 1.2 σ calculated after modelling of the phosphoserine residues is shown overlaid with the $F_o$-$F_c$ difference map (red) calculated before modelling and contoured at 2.6 σ. **b.** Same views and maps (including contour levels) for the HipBST$^{S59A}$ structure.

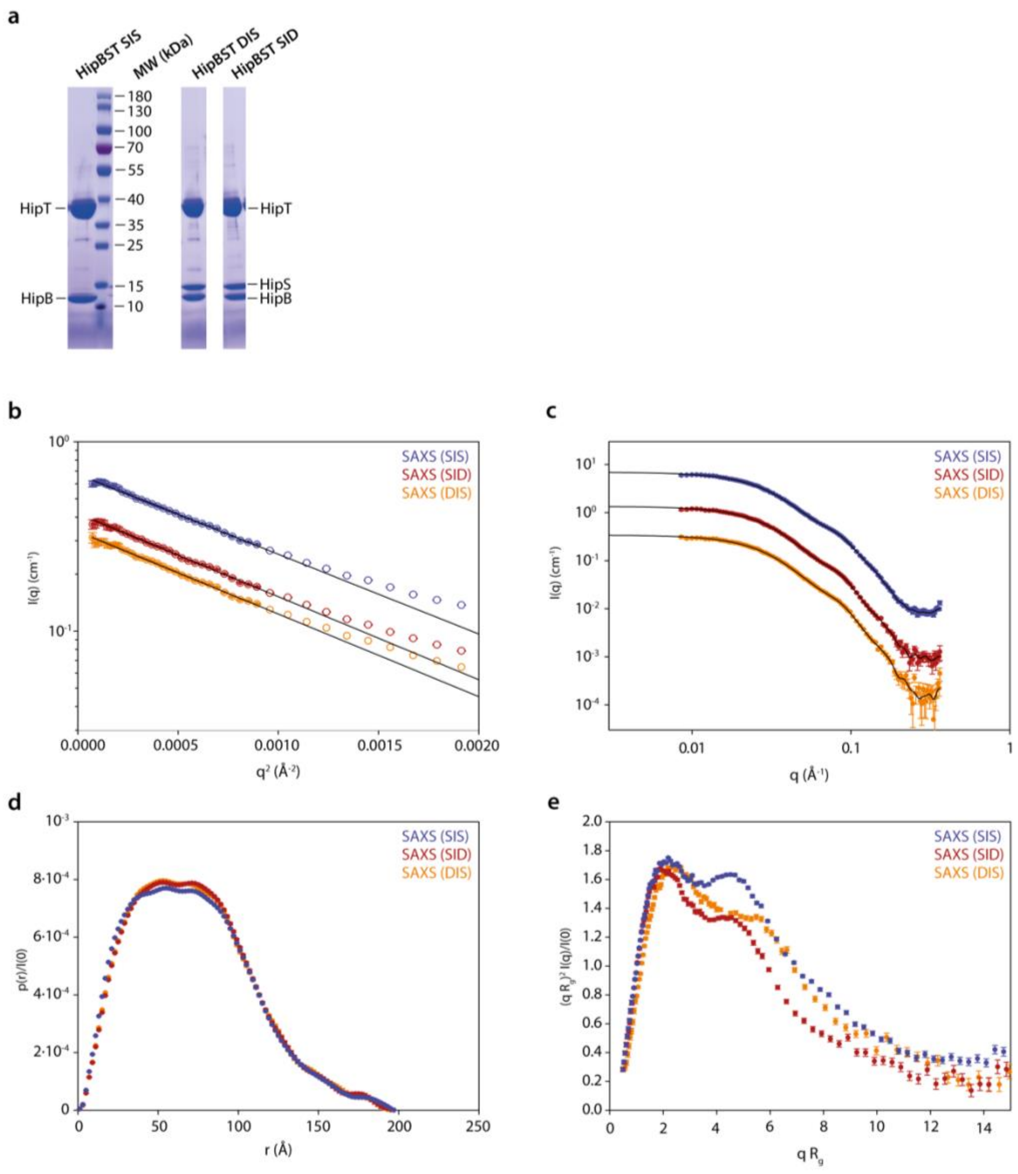


**Figure 4 supplement 1. SAXS analysis of HipBST variants. a.** SDS-PAGE gels with purified samples of HipBST (D210A, SIS), HipBST (D210A, DIS), and HipBST (D210A, SID) showing that the HipS band is missing in the SIS purification (left). **b.** Guinier plots giving radii of gyration of 54.9 ± 0.3 Å (DIS), 55.1 ± 0.3 Å (SID), and 54.0 ± 0.2 Å (SIS). **c.** Indirect Fourier transformations giving radii of gyration of 57.7 ± 0.4 Å (DIS), 57.9 ± 0.3 Å (SID), and 57.5 ± 0.3 Å (SIS). **d.** Pair distance distribution function. **e.** Normalised Kratky plot using the values of $R_g$ and $I(0)$ from the Guinier fit for normalisation.

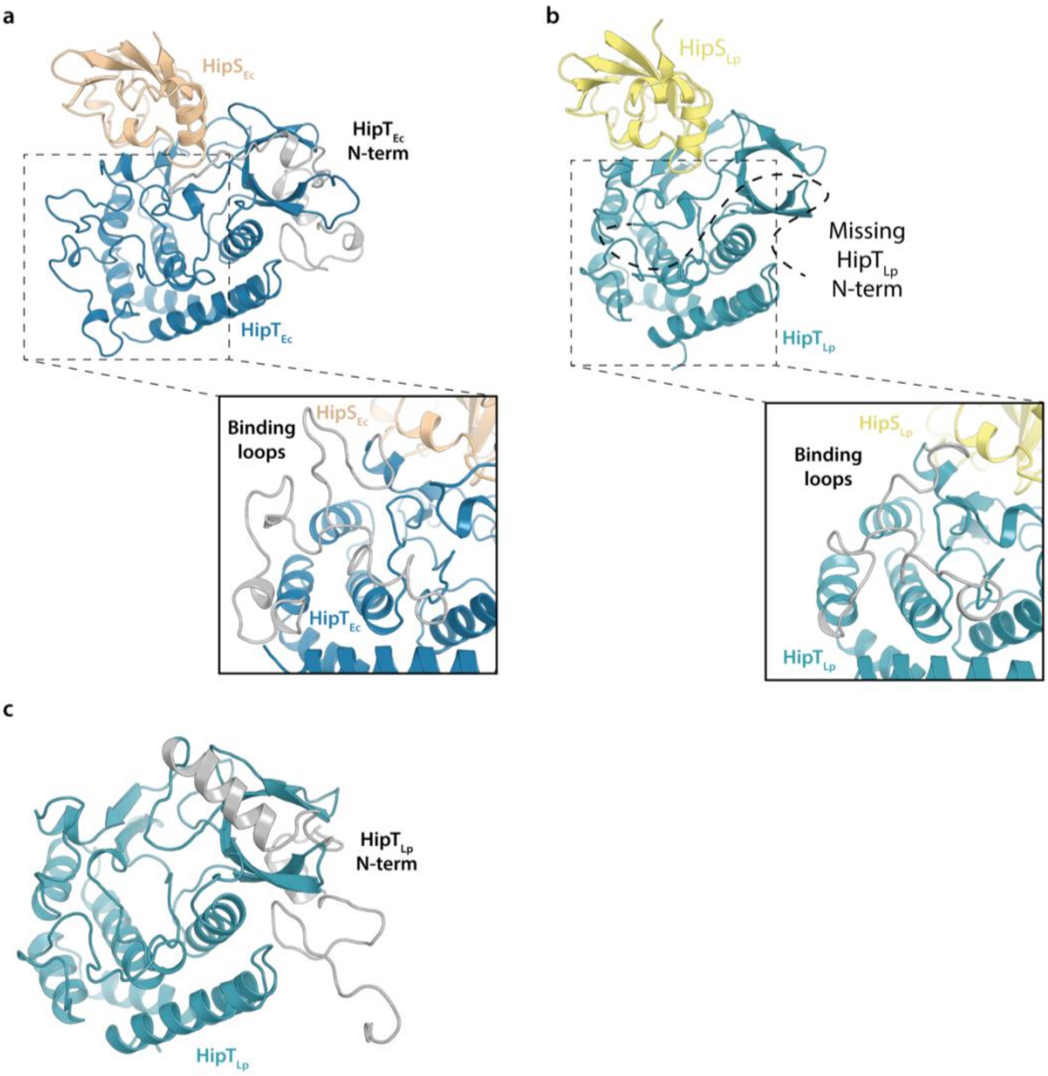


**Figure 6 supplement 1. Structural comparison between HipST$_{Ec}$ and HipST$_{Lp}$. a.** Structure of the HipST heterodimer from the HipBST$^{D233Q}$ complex showing two distinct regions, the N-terminal minidomain (grey), which is ordered in this structure and the target binding regions comprised of several loops (zoomed view) (Gerdes *et al.*, 2021). **b.** The crystal structure of HipST$_{Lp}$ (PDB: 7VKB) (Zhen *et al.*, 2022) does not have a visible N-terminal domain when HipS is bound, and the loops forming the target-binding region are different (zoomed view). **c.** The N-terminal domain of HipT$_{Lp}$ (grey) is resolved when HipS is not present.
